# An open-source, externally validated neural network algorithm to recognize daily-life gait of older adults based on the lower-back sensor

**DOI:** 10.1101/2025.01.31.635928

**Authors:** Yuge Zhang, Sjoerd M. Bruijn, Michiel Punt, Jorunn L. Helbostad, Mirjam Pijnappels, Sina David

## Abstract

**Background:** Accurate gait recognition from daily physical activities is a critical first step for further fall risk assessment and rehabilitation monitoring based on inertial sensors. However, most openly available models are based on healthy young adults ambulating in structured conditions.

**Objective:** This study aimed to develop an open-source and externally validated algorithm for daily-life gait recognition of older adults based on acceleration and angular velocity, as well as acceleration data only, and explore the effect of the use of data augmentation in the model training.

**Methods:** A convolutional neural network was trained for gait recognition. The data for model training was lower-back inertial sensor data from 20 older adults (mean age 76 years old), with annotated synchronized activity labels in semi-structured and daily-life conditions. The data was randomly split into training, validation, and testing datasets by participants, and the model was trained multiple times using these different splits. The model was trained based on data from six channels (accelerations and angular velocities) and three channels (accelerations only) under conditions with and without data augmentation, respectively. External validation was evaluated based on lower-back sensor data collected from 47 stroke survivors (mean age 72.3 years old) in balance and walking tests.

**Results:** For the testing dataset, the median accuracy ranged from 94 % to 98 %, precision from 63 % to 85 %, sensitivity from 95 % to 97 %, F1-score from 76 % to 90 %, and specificity from 94 % to 98 %. For the external validation dataset, the median accuracy ranged from 97 % to 100 %, precision from 99.9 % to 100 %, sensitivity from 71 % to 100 %, F1-score from 83 % to 100 %, and specificity 100 %.

**Conclusions:** Based on lower-back-worn inertial sensor data, we provide an accurate, open-source, and externally validated daily-life gait recognition algorithm for older adults, with one model for six-axis input data and another for three-axis input data. Besides, we found when training the model, the use of data augmentation is especially helpful on the model based on acceleration data only.

## Introduction

Each year, approximately one-third of older adults experience falls [1, 2] , most of which occur during walking [3]. Many studies have shown the potential of gait measures derived from inertial measurement units (IMUs) to predict fall risks [4, 5], and rehabilitation monitoring, such as for stroke survivors [6]. However, to extract these measures, the first crucial step is to recognize gait among various daily-life physical activities accurately is necessary and crucial.

Convolutional neural networks (CNN) can learn features via filters (or kernels) and have been increasingly used for gait recognition on IMU data [7]. Most of the developed models are based on young adults and their accuracy in recognizing gait is greater than 88 % [8, 9]. However, older adults typically walk slower or suffer from gait impairments, thus models trained on young adults may not be suitable for older adults.

Several challenges need to be recognized and overcome to apply such models to older adults. First, the data collection conditions affect the model’s robustness. Most studies used data that were collected in a structured protocol in the lab [10, 11] or in semi-structured conditions. In the semi-structure conditions, activities can be performed out of sequence, labeled by the experimenter in a simulated environment, or by participants themselves using an electronic device in a daily life condition [12-14]. However, such activities are different from daily life in context, behavior, and rhythm [15] , and especially when there is an observer on site, activities might be influenced by the Hawthorne effect [16]. Because of these challenges, unsupervised data with valid activity labels collected in daily life is lacking. One of the few exceptions is the data set of Bourke and colleagues who collected an IMU dataset (the ADAPT dataset) of older adults, including standardized data in semi-structured activities, and data in daily-life conditions [17], labeled by synchronized video footage from a body-worn camera.

Second, daily-life data is often imbalanced in activities with a smaller amount of gait. Therefore, sensitivity and F1-score are important to evaluate the gait recognition model. Moreover, to evaluate the robustness of a trained model, external validation of the model is necessary. As an example, in the study of Zou[9], the CNN model achieved an accuracy of 92.9 % for gait on their training dataset but only 40.6 % on the external dataset, showcasing the importance of testing and reporting results for external validation data. Third, to increase the robustness of models, the variability of the data should be considered. Data augmentation (DA) can expand a dataset synthetically by transforming existing samples, increasing the training data’s quantity and variability [18]. Um and colleagues found that adding rotated data can alleviate the effect of variability of sensor orientations, thereby improving accuracy by 11.9 % on the testing dataset [19].

Finally, only a few studies have made their algorithms available in an easy-to-use software repository [9, 20], which limits their accessibility, practical application, and potential impact. Therefore, to overcome all these challenges, we aimed to develop an open-source externally validated neural network algorithm for recognizing daily-life gait based on lower back IMU data of older adults. Additionally, we will explore the effects of rotation as data augmentation on the model based on 3 or 6-channel IMU data.

## Methods

### Datasets

We used two datasets: one was the ADAPT dataset for model training [17] and the second one was a dataset of stroke survivors serving as external validation [21, 22]. The data description and the code belonging to this paper (including the trained CNNs) are available at [23]. Details of the participants’ characteristics are presented in Table 1.

**Table 1.** Demographic characteristics of participants.

| Characteristics | ADAPT | Balance tests in External dataset | Walking tests in External dataset |
| --- | --- | --- | --- |
| Number | 20 | 47 | 18 |
| Age (years) (Mean (SD)) | 76.4 (5.6) | 72.3 (12.2) | 68.4 (9.3) |
| Sex (Male/Female) | 5/15 | 30/17 | 12/6 |
| Height (m) (Mean (SD)) | 1.67 (0.07) | 1.75 (0.11) | 1.74 (0.12) |
| Weight (kg) (Mean (SD)) | 73.7 (11.4) | 78.2 (14.5) | 79.7 (15.4) |
| Percent of participants with disabilities/diseases that affect activity | 60 % | 100 % | 100 % |
| Hemiparetic side (left/right/both/unknown) | N.A. | 26/10/8/3 | 8/4/5/1 |
| Stroke types (ischemic/hemorrhagic/subarachnoid) | N.A. | 38/6/3 | 14/3/1/0 |
| Gait speed (m/s) (Mean (SD)) | N.A. | 0 | 1.3 (0.6) |
| Walking bout (s) (Median (IQR)) | 4.1 (4.5) | 0 | 120 (0) |
| Percent of participants with falls in the past year | 55 % | N.A. | N.A. |

*SD: Standard deviation, IQR: Interquartile Range, the difference between the 75th and 25th percentiles of the data*

### Training dataset

The ADAPT dataset is a comprehensive physical activity reference dataset, for which 20 older adults performed various activities in semi-structured and daily-life environments, wearing 12 body-worn IMUs and a synchronized chest-mounted GoPro camera [17]. We were primarily interested in the lower-back sensor data as the clinical value of the lower back for trunk stability, balance control, and fall risk assessment [24]. The data was collected by the Usense (University of Bologna, Italy) sensor at a sampling frequency of 100 Hz. All participants were able to take verbal instructions and walk independently for 100 m without a walking aid. Twelve participants had self-reported disabilities or diseases affecting activity (unspecified). Eight participants have experienced one or more falls in the past year.

This dataset contained 43.9 hours of activities in total, including 9.5 hours in semi-structured conditions and 34.4 hours in daily-life conditions. All activities were annotated according to strict activity definitions based on the video footage [25]. There were 16 labeled activities: walking, walking with transitions, shuffling, standing, sitting, transition, leaning, jumping, dynamic, static, lying, shaking, picking, kneeling, stair ascending, and descending. Walking was defined as locomotion towards a destination with one stride or more, finishing with one foot placed beside the other foot, also including certain steps during the transition from walking to another posture [25]. There were 6.13 hours of walking in total, accounting for 14 % of all activities, and the agreement of video labels by raters was >90 % [17]. In model training, all activities not labeled as walking were categorized as non-walking. Each participant’s distribution of walking bouts is shown in Supplementary Figure 1.

### External validation dataset

The external validation data was collected by Felius and colleagues [21, 22] from 47 stroke survivors wearing an inertial sensor (Aemics b.v. Oldenzaal, The Netherlands) sampling at 104 Hz on the lower back (L5). Data from balance tests and walking tests were categorized as non-walking data and walking data, respectively. All participants could independently complete balance tests, and 18 of them could independently walk. To make it consistent with the ADAPT data, in this study, we used the data where participants walked without using aids. For completeness, we also provided the information of participants who walked with aids, shown in Supplementary Table S2.

The balance tests included one condition of sitting and four conditions of unsupported standing (eyes open, closed, feet together, and foam). The walking test was conducted on a 14-meter-long surface walkway for two minutes, including turns at both ends [22]. Participants walked at a self-selected pace without stopping, being distracted, or developing observable clonus. As a calibration, participants remained stationary before and after the tests for a period.

### Data processing

Figure 1 gives an overview of the data processing for model training and evaluation. For model training, the steps included data preparation, splitting, augmentation, segmentation, and balancing. For the external validation, steps included data preparation, augmentation, and segmentation. We used the input of six channels (acceleration and angular velocity) and three channels (acceleration only) separately. Matlab 2021 and Python (version 3.8) were used.

**Figure 1.**
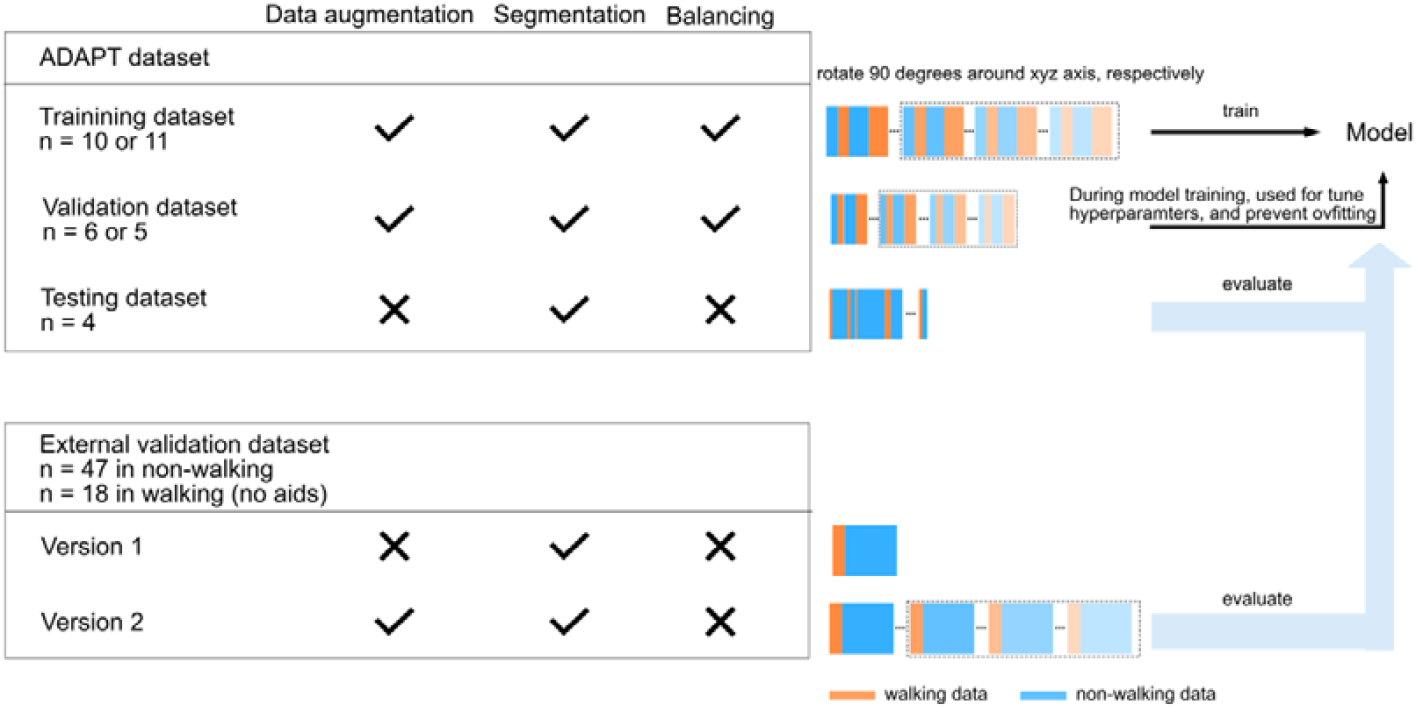
The data process for model training and evaluation.

### Data preparation

The ADAPT dataset was organized as six-channel IMU data, activities labels, and the participant’s ID, resulting in a 16007994 x 6 input matrix, a 16007994 x 1 target matrix, and a 16007994 x 1 group matrix, respectively. For the external dataset, after deleting the calibration data, the dataset was organized as IMU data, labels, and participants’ ID, resulting in a 2523115 x 6 input matrix, 2523115 x 1 target matrix, and a 2523115 x 1 group matrix, respectively.

### Data splitting

We used a repeated hold-out validation method to split the ADAPT dataset for model training. Each time, the dataset was randomly divided by participants, resulting in the percentage of training, validation, and testing datasets as 55%, 25 and 20%.

### Data augmentation (DA)

The training and validation datasets from ADAPT were augmented by applying 90° rotation to each signal channel to simulate major sensor placement shifts. For comparison, we also trained the model without data augmentation. To increase the sample size with different sensor orientations, the external dataset was augmented as well.

### Data segmentation

Studies indicate that smaller windows were preferable for low-intensity activities and older adults [26]. An interval of 1-2 seconds seems the best trade-off between accuracy and recognition time [27, 28]. Therefore, before being fed into the model, all datasets including the external validation data were segmented into 2-second windows with 50 % overlap, with each window assigned a single activity label. Additionally, in each window, data was normalized by subtracting the corresponding mean value on each axis.

### Data balancing

The dataset for model training was imbalanced, containing 15235 walking windows and 131208 non-walking windows (a rate of 1:8.61). Such imbalances can bias model performance [32, 33]. To address this, for both the training and validating datasets, we down-sampled the number of non-walking windows to the same as the number of walking windows. The testing and external validation datasets were not down-sampled to maintain the uncertainty and diversity of data. The number of walking windows in the external validation dataset was 11271 and the number of non-walking windows was 89139, indicating a rate of 1:7.91.

### CNN Model Structure

The structure of the CNN model developed in this study contains the following layers.

1. An input layer. The partitioned signal data fed into the first layer was a 3D matrix form of (batch size × windows × number of channels), where batch size B = 32, sliding window size W = 200 (which resembles 2 seconds), and the number of channels N = 6 (i.e. acceleration and angular velocity data), or N = 3 (i.e. acceleration only).
2. Hidden layers, including two convolutional layers, one max pooling layer, and one fully connected layer. A convolution layer with multiple filters creates multiple feature maps, enabling the extraction of a diverse set of features from the input [29]. Here, we used 64 filters, so that each convolutional layer has 64 feature maps from the previous layer. The weights of the filters are automatically updated in subsequent training iterations (epochs). The max pooling layer reduces model parameters by maintaining the maximum values of each feature map. The fully connected layer converts the multidimensional array from the previous layer to a single-dimensional vector.
3. Dropout layers. Although the representational ability of deep neural networks is compelling, overfitting remains a significant drawback due to the non-linear hidden layers [30, 31]. By randomly setting the output of a portion of the neurons to zero, using dropout in the training process reduces the neuron dependencies and significantly improves the generalization ability. In our study, a dropout layer with a rate of 50 % was added after each convolutional and pooling layer.
4. The output layer. This layer leverages features from the fully connected layer to generate the probability distribution of the categories, generally adopting the ‘Softmax’ function.

To prevent the network from overfitting, we also used ‘Early Stopping’ to stop training when the loss value did not improve for 3 consecutive epochs.

To evaluate model performance, we calculated the accuracy, precision, sensitivity, F1-score, and specificity between predicted labels and true labels for the validation, testing, and external validation datasets.

## Results

The best-trained models of each channel can be found in easy-to-use functions [23], which were from 6 channels without DA and 3 channels with DA, respectively, shown in Table 3.

**Table 3.** Summary of gait recognition performance of the best models.

| Training model on 6 channels without DA |  |  |  |  |  |
| --- | --- | --- | --- | --- | --- |
| Dataset | Accuracy | Precision | Sensitivity | F1-score | Specificity |
| Validation | 94.2 % | 90.6 % | 98.7 % | 94.4 % | 89.8 % |
| Testing | 93.3 % | 60.6 % | 99.2 % | 75.3 % | 92.7 % |
| External validation with DA | 99.8 % | 98.8 % | 99.5 % | 99.2 % | 99.9 % |
| External validation without DA | 100.0 % | 99.8 % | 100.0 % | 99.9 % | 100.0 % |
| Training model on 3 channels with DA |  |  |  |  |  |
| Dataset | Accuracy | Precision | Sensitivity | F1-score | Specificity |
| Validation | 96.4 % | 95.6 % | 97.2 % | 96.4 % | 95.5 % |
| Testing | 96.5 % | 78.7 % | 98.9 % | 87.6 % | 96.1 % |
| External validation with DA | 99.9 % | 99.7 % | 99.2 % | 99.4 % | 100.0 % |
| External validation without DA | 99.9 % | 99.9 % | 99.6 % | 99.7 % | 100.0 % |

After training our models using different split datasets, we obtained the median (interquartile range, IQR) of model performance and the best model for each number of channels, the results of which are shown in Figure 2. Detailed values are shown in Supplementary Table S1, which also includes results of the external datasets using walking aids.

**Figure 2.**
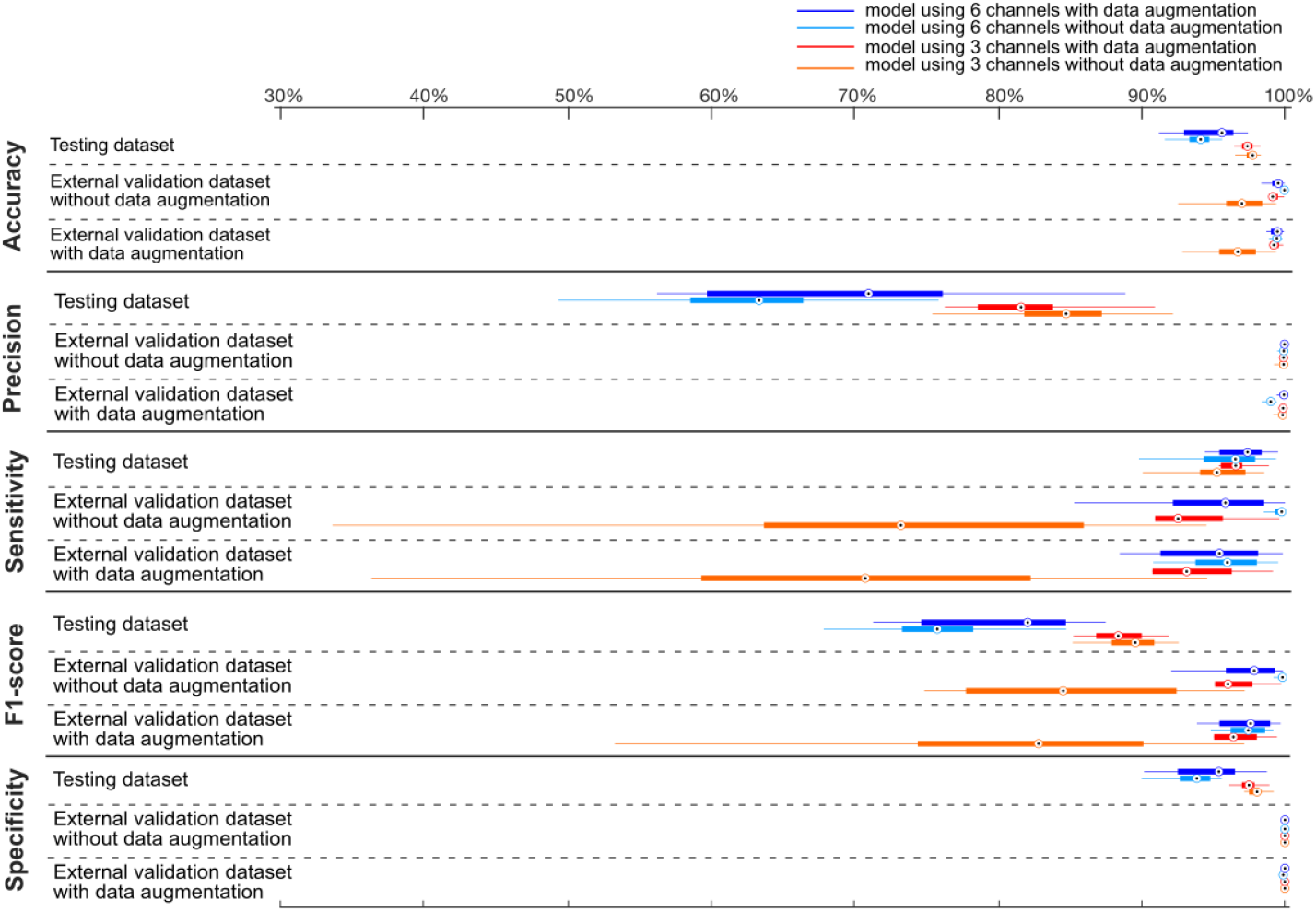
The median (IQR) of model performance over multiple training times. *(DA: data augmentation; IQR: the Interquartile Range, the difference between the 75th and 25th percentiles of the data)*.

For the testing dataset, the range of median values of performance across all types of models were [94.1 %, 97.8 %] for accuracy, [63.4 %, 84.8 %] for precision, [95.3 %, 97.4 %] for sensitivity, [75.8 %, 89.6 %] for F1-scores and [93.9 %, 98.1 %] for specificity. For the testing dataset, the 3-channel model performed overall better than the 6-channel model, no matter whether using DA or not. In addition, using DA in the 6-channel model increased the precision and F1-score by 7.6 % and 6.3 %, while having little influence on its accuracy, sensitivity, and specificity.

For the external validation dataset, the range of median values of performance across all types of models was [97 %, 100 %] for accuracy, [99.9 - 100 %] for precision, [73 %, 100 %] for sensitivity, [85 %, 100 %] for f1-scores, and 100 % for specificity. In contrast to the testing dataset, in the external validation dataset, the 6-channel model performed generally better than the 3-channel model. In addition, using DA had little influence on the 6-channel model but greatly increased the median sensitivity on the 3-channel model i.e., from 70. 8 % to 93.2 % in external validation data with augmentation, from 73.2 % to 92.5 % in the raw external validation data. Also, it increased the median F1-score on the 3-channel model, by 13.6 % and 11.5 % on external validation data with and without DA, respectively.

Furthermore, in Supplementary Table S1, we showed the performance on the external validation for the participants who used walking aids. We found the 6-channel model performed still well when walking with aids, as the median sensitivity ranged from 73 % to 89 %. However, the 3-channel model without data augmentation performed worst with a median range of sensitivity of 48 %, after using data augmentation, the value greatly increased to 66 %.

## Discussion

We developed an open-source, externally validated CNN model to recognize daily-life gait in older adults and explored the effect of the use of data augmentation in model training. For the model with 6-channel input data (both acceleration and angular velocity), the overall accuracy exceeded 93.3 %, with precision over 60.6 %, sensitivity over 98.7 %, F1-score over 75.3 %, and specificity over 90 % across testing and external validation datasets. For the model with 3-channel input data (acceleration only), the overall accuracy was over 96.4%, with precision over 78.7 %, sensitivity over 97.2 %, F1 score over 87.6 %, and specificity over 95.5%.

In the testing dataset, the 3-channel model performed better than the 6-channel model in accuracy, precision, F1-score, and specificity, except for the sensitivity, which was similar. This was not consistent with other studies that showed that adding angular velocities can be complementary to increasing the model’s sensitivity [9, 32-34]. This could be caused by overfitting in the 3-channel data, although during the model training, we used early stopping to halt the process before the model over-specialized on the training data. Adding a larger sample size to the training dataset may decrease the likelihood of overfitting. In addition, data augmentation had little influence on training the 3-channel model (median accuracy 97.4 % vs 97.8%, precision 81.6 % vs 84.8 %, sensitivity 96.6 % vs 95.3 %, F1-score 88.4 % vs 89.6 %, specificity 97.5 % vs 98.1 %, for with and without DA), but it can increase the precision and F1-score in the 6-channel model (71.0 % vs 63.4 %, 82.1 % vs 75.8 % for with and without DA). This indicates that such data augmentation may prove to be especially important when only accelerometer data (as opposed to accelerometer and gyroscope data) is used, as accelerometer data alone may not capture gait fully.

In the external validation dataset, the accuracy of models was greater than 94 %, which is a huge improvement when compared to the 40.6 % reported by Zou et al [9] on the external validation data collected in structural conditions. Additionally, the 6-channel model performed better than the 3-channel model, which is in line with other studies [9, 32-34]. Using data augmentation had little influence on the 6-channel model but greatly increased the median sensitivity and F1-score of the 3-channel model (in the 6-channel model, 95.8 % vs 99.8 %, 97.9 % vs 99.8 %, for with and without DA; in the 3-channel model, 92.5 % vs 73.2 %, 96.0 % vs 84.5 %, for with and without DA). This suggests that data augmentation indeed had added value for 3-channel data.

Although both testing and external validation datasets were imbalanced, there were three main differences between them, which may cause the different performances. Firstly, the testing dataset was collected in daily-life conditions and the external data was collected in a structured condition. Due to this, in the testing dataset, for each participant’s data, gait was alternated with other activities, such as stand-walk-sit and sit-walk-stairs. However, in the external validation dataset, because gait and non-gait were dispersed in data collection, the same activity was pooled across all subjects. Meanwhile, the daily-life environment may also alter gait patterns, e.g., slopes and obstacles may affect how people walk, and people may walk differently indoors and outdoors [24]. Therefore, activity context may significantly influence the model performance [15], as evidenced by the overall better performance on the structured external dataset than on the daily-life testing dataset. On the other hand, this also indicates that a model trained on daily-life data can also perform well on structured data.

Secondly, the testing dataset had various non-gait activities, such as shuffling, walking with transitions, and going up and down the stairs. The external dataset included only two static activities, which were sitting and standing. Studies showed that the sensor data of dynamic activities, such as going up and down stairs may be more easily misclassified as gait [35, 36]. Therefore, the non-gait activities in the testing dataset were easier to distinguish so the model performance was better.

Lastly, studies showed that a short walking duration and slow walking speed negatively affect the model performances [24, 37]. In the testing dataset, as shown in the supplementary Figure S1, most walking bouts were between 2s and 6.45 seconds (median+IQR/2), and the longest walking bouts were shorter than 170 seconds. Because parts of these bouts are needed to accelerate and decelerate [38], the walking speed of the testing dataset can be expected to be relatively slower than observed in the external dataset. On the contrary, in the external data, even though participants were stroke survivors, they walked independently with an average speed of 1.3 m/s in a 2-minute walkway (calculated from the IMU data) [22]. Such a situation may also explain why models performed better on the external validation dataset than the testing dataset. Van Ancum demonstrated that the 4-m gait speed is only related to the high end of daily-life speed and cannot reflect the walking ability of daily life [39]. Therefore, further studies should collect more daily-life data indoors and outdoors for training and external validation, to cover various walking speeds, walking distances, and contexts [34].

The results on walking using aids showed similar results as for walking without aids, even though our models were trained on only walking without aids, suggesting that our models are applicable to a wide range of gaits.

## Strengths and Limitations

The model we developed can correctly recognize the daily-life gait of older adults and was externally validated on stroke survivors. In addition, we provided the model for different input channels and explored the effect of the use of data augmentation in the model training. Last but not least, we offer the best models and code for step-by-step data processing to obtain gait episodes from either 3 or 6-channel data. Users can directly use and refine our models from the GitHub repository [23].

There are also some limitations of our study. Firstly, the external dataset was structured, which may have inflated model performance. Thus, model performance on an external daily-life dataset is unclear yet. Secondly, we used down-sampling methods to balance our training data, thereby reducing the number of available training windows. More importantly, the training data in our study consisted of only 10 or 11 participants, insufficient to cover the varied gait patterns of a wide population of older adults, thus, increasing the number of participants for training is recommended to improve the robustness. Lastly, in this study, we used 50 % overlapping windows. Further studies can try to use an overlap of 95 %-99 % and then use approaches like majority voting machine learning for a more accurate model prediction.

## Conclusions

In conclusion, this study provided a good convolutional neural network for daily-life gait recognition based on lower-back IMU data for healthy and gait-impaired older adults, available on (https://github.com/ZYuge/Recognize-real-world-walking-based-on-deep-learning-methods). Besides, we found when training the model, the use of data augmentation is especially helpful on the model based on acceleration data only. The open-source code promises to promote community-based remote gait monitoring.

## Abbreviations

IMU: inertial measurement units
CNN: convolutional neural networks
ADAPT: a personalized fall risk assessment system for promoting independent living

## Acknowledgments

For the use of the ADAPT data, we are grateful to the NextMove Laboratory at the Department of Neuroscience in the Faculty of Medicine at the Norwegian University of Science and Technology in Trondheim, Norway. The ADAPT project was led by Professor Jorunn L. Helbostad and Dr. Espen Alexander F. Ihlen, the data was handled and provided by Dr. Sabato Melone and was funded by the Norwegian Research Council.

We also would like to thank all team members of Richard A.W. Felius for providing external validation data, at the Research Group Lifestyle and Health, Utrecht University of Applied Sciences, Utrecht, the Netherlands.

The conceptualization and design of the study: YZ, SMB, MPij; Data collection: JLH, MPu; Data analysis: YZ, SD, SMB, MPu; Interpretation of results: YZ, SD, SMB, MPu, MPij; Write and revise draft: YZ, SD, SMB, MPu, JLH, MPij; Supervision: SD, SMB, MPij. YZ was funded by the China Scholarship Council(CSC) (202009110145). MPij and SMB were funded by a grant from the Dutch Organization for Scientific Research (NWO) (no. 91714344 and 016.Vidi.178.014, respectively).

