## Supplementary for "An open-source, externally validated neural network algorithm to recognize daily-life gait of older adults based on the lower-back sensor"

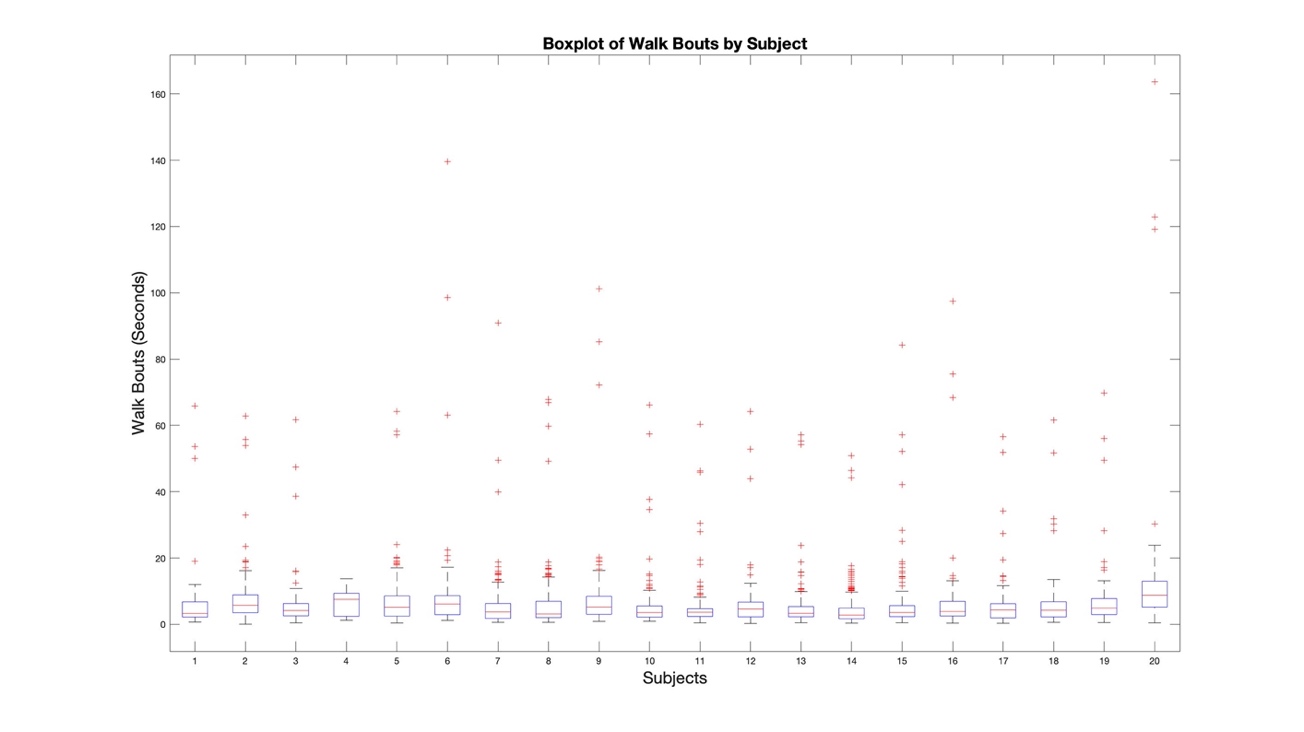

**Figure S1.** Boxplot of the time of walking bouts for each of the participants in the ADAPT dataset. The top edge (Q3), the middle line(Q2), and the bottom edge (Q1) of the box represent the 75^th^, 50^th,^ and 25^th^ percentile, respectively. The upper whisker (Q4) represents the maximum, 1.5 times the interquartile range above Q3. The lower whisker represents the minimum, 1.5 times the interquartile range below Q1. Data points that fall outside the range of the whiskers are marked as outliers.

**Table S1.** The performance gait classification based on the 4 datasets of either 3 or 6 channels and the with and without data augmentation (DA) (Median (IQR), unit: %)(responding to the boxplot)

| **Models** | **Dataset** | **DA(yes/no)** | **Accuracy** | **Precision** | **Sensitivity** | **F1-score** | **Specificity** |
| --- | --- | --- | --- | --- | --- | --- | --- |
| 6 channels, with DA | Testing | no | 95.6 (3.4) | 71.0 (16.4) | 97.4 (2.9) | 82.1 (10.1) | 95.4 (4.0) |
| 6 channels, without DA | Testing | no | 94.1 (1.4) | 63.4 (7.9) | 96.5 (3.6) | 75.8 (5.0) | 93.9 (2.1) |
| 3 channels, with DA | Testing | no | 97.4 (0.7) | 81.6 (5.2) | 96.6 (1.5) | 88.4 (3.2) | 97.5 (0.9) |
| 3 channels, without DA | Testing | no | 97.8 (0.6) | 84.8 (5.4) | 95.3 (3.2) | 89.6 (3.0) | 98.1 (0.7) |
| **Range** |  |  | **[94.1,97.8]** | **[63.4,84.8]** | **[95.3,97.4]** | **[75.8,89.6]** | **[93.9,98.1]** |
| **No Aids in walking** |  |  |  |  |  |  |  |
| 6 channels, with DA | Ex-validation | no | 99.5 (0.7) | 100.0 (0.1) | 95.8 (6.4) | 97.9 (3.4) | 100.0 (0.0) |
| 6 channels, without DA | Ex-validation | no | 100.0 (0.1) | 99.9 (0.2) | 99.8 (0.7) | 99.8 (0.3) | 100.0 (0.0) |
| 3 channels, with DA | Ex-validation | no | 99.1 (0.5) | 99.9 (0.1) | 92.5 (4.7) | 96.0 (2.6) | 100.0 (0.0) |
| 3 channels, without DA | Ex-validation | no | 97.0 (2.5) | 99.9 (0.4) | 73.2 (22.3) | 84.5 (14.7) | 100.0 (0.0) |
| **Range** |  |  | **[97,100]** | **[99.9,100]** | **[73.2,99.8]** | **[84.5,99.8]** | **[100,100]** |
| 6 channels. with DA | Ext-validation | yes | 99.5 (0.8) | 99.9 (0.3) | 95.4 (6.8) | 97.6 (3.5) | 100.0 (0.0) |
| 6 channels, without DA | Ext-validation | yes | 99.4 (0.5) | 99.0 (0.3) | 96.0 (4.3) | 97.4 (2.4) | 99.9 (0.0) |
| 3 channels, with DA | Ext-validation | yes | 99.2 (0.6) | 99.9 (0.2) | 93.2 (5.5) | 96.4 (3.0) | 100.0 (0.0) |
| 3 channels, without DA | Ext-validation | yes | 96.7 (2.5) | 99.8 (0.3) | 70.8 (22.9) | 82.8 (15.7) | 100.0 (0.0) |
| **Range** |  |  | **[96.7,99.5]** | **[99,99.9]** | **[70.8,96]** | **[82.8,97.6]** | **[99.9,100]** |
| **With Aids in walking** |  |  |  |  |  |  |  |
| 6 channels, with DA | Ex-validation | no | 94.2 (7.2) | 100 (0.1) | 72.6 (33.7) | 84.1 (22.3) | 100 (0) |
| 6 channels. without DA | Ex-validation | no | 97.7 (0.9) | 99.9 (0.1) | 89.2 (4.2) | 94.3 (2.3) | 100 (0) |
| 3 channels, with DA | Ex-validation | no | 92.8 (1.5) | 99.9 (0.1) | 66 (6.9) | 79.5 (4.9) | 100 (0) |
| 3 channels, without DA | Ex-validation | no | 88.9 (3.3) | 99.9 (0.2) | 48.1 (15.3) | 64.9 (13.7) | 100 (0) |
| **Range** |  |  | [88.9,97.7] | [99.9,100] | [48.1,89.2] | [64.9,94.3] | [100,100] |
| 6 channels, with DA | Ext-validation | yes | 93.7 (5.5) | 99.9 (0.1) | 70.2 (26.3) | 82.4 (16.9) | 100 (0) |
| 6 channels, without DA | Ext-validation | yes | 96.7 (1.7) | 99.5 (0.2) | 85.1 (8.1) | 91.7 (4.6) | 99.9 (0) |
| 3 channels, with DA | Ext-validation | yes | 94.3 (1.4) | 99.9 (0.1) | 73 (6.6) | 84.4 (4.4) | 100 (0) |
| 3 channels, without DA | Ext-validation | yes | 89.4 (2.1) | 99.9 (0.2) | 50.2 (10.1) | 66.8 (8.7) | 100 (0) |
| **Range** |  |  | [89.4,96.7] | [99.5,99.9] | [50.2,85.1] | [66.8,91.7] | [99.9,100] |

**Table S2.** Participants characteristics of the external validation data

| Characteristics | Balance tests | Walking tests (without aids) | Walking tests (with aids) |
| --- | --- | --- | --- |
| Number | 47 | 18 | 29 |
| Age (years) (Mean (SD)) | 72.3 (12.2) | 68.4(9.3) | 73.6(13) |
| Sex (Male/Female) | 30/17 | 12/6 | 19/10 |
| Height (m) (Mean (SD)) | 1.75 (0.11) | 1.74(0.12) | 1.75(0.12) |
| Weight (kg) (Mean (SD)) | 78.2 (14.5) | 79.7(15.4) | 79.0(14.7) |
| Percent of participants with disabilities/diseases that affect activity | 100% | 100% | 100% |
| Hemiparetic side (left/right/both/unknown) | 26/10/8/3 | 8/4/5/1 | 14/8/5/2 |
| Stroke types (ischemic/hemorrhagic/ subarachnoid) | 38/6/3 | 14/3/1/0 | 26/2/1 |
| Gait speed (m/s) (Mean (SD)) | 0 | 1.3 (0.6) | 1.3 (0.6) |
| Walking bout (s) (Median(IQR) | 0 | 120 (0) | 120 (0) |
| Percent of participants with falls in the past year | N.A. | N.A. | N.A. |
